# Clonal plasma cells in AL amyloidosis are dependent on pro-survival BCL-2 family proteins and sensitive to BH3 mimetics

**DOI:** 10.1101/542159

**Authors:** Cameron Fraser, Adam Presser, Vaishali Sanchorawala, Shayna R. Sarosiek, Kristopher A. Sarosiek

**Affiliations:** John B. Little Center for Radiation Sciences, Department of Environmental Health, Harvard T.H. Chan School of Public Health, Boston MA; Laboratory of Systems Pharmacology, Harvard Medical School, Boston MA; Section of Hematology & Medical Oncology, Boston Medical Center, Boston MA; Amlyloidosis Center, Boston University School of Medicine, Boston MA

**Keywords:** BH3 mimetics, AL amyloidosis, BCL-2, MCL-1, bortezomib

## Abstract

Immunoglobulin light chain (AL) amyloidosis is a protein misfolding disorder characterized by the production of amyloidogenic immunoglobulin light chains by clonal populations of plasma cells. These abnormal light chains misfold and accumulate as amyloid fibrils in healthy tissues causing devastating multi-organ dysfunction that is rapidly fatal. Current treatment regimens, which include proteasome inhibitors, alkylating agents, and immunomodulatory agents, were developed for the treatment of the more common plasma cell disease, multiple myeloma, and have limited efficacy in AL amyloidosis as demonstrated by the median survival of 2-3 years. The recent development of novel small-molecule inhibitors of the major pro-survival proteins from the apoptosis-regulating BCL-2 family has created an opportunity to therapeutically target abnormal cell populations, yet identifying the extent of these dependencies and how to target them clinically has thus far been challenging. Using bone marrow-derived plasma cells from 45 patients with AL amyloidosis, we find that clonal plasma cells are highly primed to undergo apoptosis and exhibit strong dependencies on pro-survival BCL-2 family proteins. Specifically, we find that clonal plasma cells in a majority of patients are highly dependent on the pro-survival protein MCL-1 and undergo apoptosis when treated with an MCL-1 inhibitor as a single agent. In addition, BCL-2 inhibition sensitizes clonal plasma cells to several current standard of care therapies. Our results suggest that BH3 mimetics, when deployed rationally, may be highly effective therapies for AL amyloidosis.

## Introduction

The systemic amyloidoses are a diverse group of disorders characterized by the abnormal deposition of protein fibrils in tissues, causing organ damage and dysfunction^1–3^. Within this group of diseases, immunoglobulin light chain (AL) amyloidosis is the most common, accounting for 70% of all diagnoses^4,5^. In AL amyloidosis, an abnormal clonal population of plasma or lymphoplasmacytic cells, typically occupying less than 10% of the bone marrow, produces monoclonal kappa or lambda light chains^6–9^. The monoclonal light chains misfold and aggregate in tissues as insoluble amyloid fibrils, ultimately causing multi-organ dysfunction that is rapidly fatal^2,8,10,11^. Due to the relative rarity of this disorder and lack of awareness, it is often diagnosed at late stages with advanced organ involvement, resulting in a median overall survival [OS] of 2-3 years from diagnosis. Importantly, although AL amyloidosis is considered a rare disease with a historic incidence of 10 cases per million individuals per year in developed countries, the prevalence of AL amyloidosis continues to increase as diagnostic tools and clinical awareness improves^12,13^.

The complexity of AL amyloidosis, due to the unique coexistence of a clonal plasma cell disorder and the amyloid fibril-induced organ dysfunction, makes treatment of this disease challenging. Due to the potential high degree of organ damage at presentation, patients are at high risk of death and are extremely susceptible to treatment toxicity in the first few months after diagnosis^1,2,8,14–16^. Therapy for AL amyloidosis is largely focused on elimination of the clonal plasma cell population that produces the amyloidogenic light chains via risk-adapted treatment approaches based on the severity of organ involvement and the performance status of each individual. For low-risk patients (approximately 25% of those diagnosed) treatment with high-dose melphalan followed by autologous stem cell transplantation (HDM/SCT) is recommended and can achieve hematologic complete responses in 25-67% of patients, organ responses in 27-83% of patients, and a median OS of 7.6 years, with some patients achieving a complete hematologic response and a median overall survival of more than 10 years^1,8,17^. The remainder of patients have intermediate or high-risk disease that is treated with a combination of therapies including the proteasome inhibitors bortezomib and ixazomib, the alkylating agents melphalan or cyclophosphamide, and glucocorticosteroids^1,2,18,19^. Various combinations of these agents, such as CyBorD or BMDex, have been evaluated clinically and treatment choices are based on expected toxicities. Although multidrug regimens are preferred, often the highest risk patients can tolerate only low-dose or single agent therapy which is unlikely to control clonal plasma cells sufficiently, leading to a median survival of only 3 to 7 months^8,20^. When making treatment decisions, consideration must be given to the individual’s performance status, severity of cardiac involvement, and risk of treatment toxicity, but also to the potential for underlying treatment resistance in the clonal plasma cells. Combination regimens may overcome typical markers of resistance, such as gain 1q21 which results in a worse outcome with oral melphalan or translocation (11;14) that may portend resistance to bortezomib^21,22^. Despite initial responses to therapy, most patients experience a relapse in disease that results in significant morbidity or mortality, therefore it is of vital importance to develop more effective and better-tolerated therapies, especially agents that may exploit inherent plasma cell vulnerabilities such as their regulation of the BCL-2 family of proteins that control apoptosis.

Apoptosis is a highly regulated form of cell death that is essential for normal mammalian development, as well as the removal of damaged, dysfunctional, or superfluous cells. The balance between pro-apoptotic and pro-survival proteins from the BCL-2 family determines whether a cell will commit to apoptosis^23–25^. When effective, most anti-cancer therapies induce an apoptotic cell death in neoplastic cells^25–27^ and we have previously found that the state of the apoptosis pathway in many types of cancers is a major determinant of their sensitivity to anti-cancer therapies^24^. Furthermore, suppression of apoptotic priming is a major driver of therapy resistance and tumor replapse^24,28–30^, strongly suggesting that apoptosis is an attractive therapeutic target.

The mitochondrial apoptosis pathway is triggered when a pro-apoptotic protein (BAX or BAK) is activated by a BH3-only protein (BIM or BID) to trigger mitochondrial outer membrane permeabilization (MOMP) and consequent release of cytochrome *c* from mitochondria to activate caspases^31,32^. However, pro-survival proteins in this family (predominantly BCL-2, BCL-X_L_, and MCL-1) can bind and block the activity of either the pro-apoptotic, pore-forming proteins (BAX or BAK) or pro-apoptotic BH3-only activators (BIM, BID, PUMA). In order for apoptosis to occur, the pro-survival proteins must be overwhelmed and BAX/BAK activated.

Since the pretreatment state of the apoptosis pathway can strongly impact therapy responses, an assay was developed that can directly measure this property: BH3 profiling. BH3 profiling functionally measures: 1) how close a cell is to the threshold of apoptosis (apoptotic priming) and 2) how dependent a cell is on specific pro-survival proteins within the BCL-2 family (apoptotic dependencies). The BH3 profiling assay is based on measuring the extent of MOMP (cytochrome c loss) in response to pro-apoptotic BH3 peptides, which mimic the activity of full-length pro-apoptotic BH3-only proteins from the BCL-2 family^33^. The extent of MOMP induced by either highly specific or indiscriminate pro-apoptotic peptides allows for the functional measurement of overall apoptotic priming as well as dependence on specific pro-survival proteins (those that can be targeted by BH3 mimetics). Notably, BH3 profiling has been used successfully to identify and target apoptotic dependencies in several human cancers. For instance, chronic lymphocytic leukemia (CLL) was originally identified as a BCL-2-dependent disease by BH3 profiling^34,35^, leading to the recent FDA approval of the specific BCL-2 inhibitor venetoclax in this malignancy^36^.

The recent development of novel small-molecule inhibitors of the major pro-survival proteins from the apoptosis-regulating BCL-2 family, called “BH3 mimetics” has created an opportunity to therapeutically eliminate cell populations that are dependent on these proteins for survival. BH3 mimetics, which can selectively target individual pro-survival proteins (e.g. ABT-199 [venetoclax] inhibits only BCL-2) or several proteins (e.g. ABT-263 [navitoclax] inhibits BCL-2, BCL-w and BCL-X_L_) and thus block the pathways that keep abnormal cells alive. These agents have been found to be highly active in hematologic malignancies. For example, venetoclax (ABT-199) has received FDA approval for treatment of chronic lymphocytic leukemia and acute myeloid leukemia based on its excellent clinical activity and tolerability^36–38^. Agents targeting the two other major pro-survival proteins (BCL-X_L_ inhibited by WEHI-539, BCL-2 and BCL-X_L_ inhibited by ABT-263, and MCL-1 inhibited by S63845) are also undergoing clinical evaluation^39–41^. Due to this favorable activity, BH3 mimetics are currently being explored in other diseases^42–44^ and in combination with other therapies^45^.

Previous data have made it clear, however, that heterogeneity in apoptotic dependencies is common within and between cancer types. Recent studies of multiple myeloma have identified clonal plasma cell vulnerabilities to BH3 mimetics, although the malignant plasma cells from patients with multiple myeloma display heterogeneity in their dependence on particular pro-survival proteins. The clonal cells have been most sensitive ex vivo to inhibition of BCL-2 and MCL-1 at initial diagnosis, with increased MCL-1 dependence in relapsed disease^46–48^. Although multiple myeloma and AL amyloidosis are related plasma cell disorders, the extent of apoptotic dependencies in AL amyloidosis has not been previously explored.

## Results

To elucidate the potential for using BH3 mimetics therapeutically in AL amyloidosis, we first sought to better understand how apoptosis is regulated in clonal plasma cells from patients that have been diagnosed with this disorder. We collected bone marrow aspirates from 45 patients with biopsy-proven AL amyloidosis that was either treatment-naïve (8 patients) or relapsed (37 patients) (Supplemental Table 1). Patients with relapsed disease had been previously treated with standard agents including proteasome inhibitors (bortezomib or ixazomib), glucocorticosteroids (dexamethasone), alkylating agents (cyclophosphamide or melphalan), anti-CD20 antibodies (rituximab), anti-CD38 antibodies (daratumamab), and/or immunomodulatory agents (pomalidomide or lenalidomide). Sixteen patients had previously undergone high dose melphalan and stem cell transplantation. At time of analysis, all 45 patients had evidence of persistent disease as determined by abnormal serum free light chain assay or serum or urine immunofixation electrophoreses. Patients were further divided by immunoglobulin light chain predominance (lambda (λ) - 37 cases; kappa (κ) - 8 cases).

After collection, bone marrow aspirates were subjected to ficoll-paque separation to isolate mononuclear cells (Supplemental Figure 1). After mononuclear cell isolation, a portion of the cells were taken for analysis by BH3 Profiling to measure 1) mitochondrial apoptotic priming and 2) extent of cellular dependence on pro-survival BCL-2 family proteins. The mononuclear cells were stained for CD38 and CD138, permeabilized, and exposed to titrated doses of pro-death signaling peptides that target the intrinsic apoptosis pathway. After exposure to the peptides was terminated, to detect the extent of mitochondrial permeabilization induced by pro-apoptotic peptides the cells were stained for cytochrome c and analyzed via flow cytometry. One patient sample was omitted from the analysis due to low cell count. The BH3 profiling results allowed us to make several key findings. First, we found that all (44 of 44) patient samples exhibited loss of cytochrome c in response to relatively high doses of pro-apoptotic BIM BH3 peptides (10 µM) (Figure 1A), which can inhibit all of the pro-survival BCL-2 family proteins^50^ and directly activate BAX^51^ or, to a lesser degree, BAK^51^. This indicates that the cells express sufficient levels of BAX and/or BAK to activate the intrinsic apoptosis pathway. Furthermore, the majority of samples exhibited loss of cytochrome c in response to even mild doses of BIM or BID BH3 (1 µM), suggesting that the plasma cells in this disorder are highly primed for apoptosis and would likely be dependent on pro-survival (anti-apoptotic) proteins to maintain their survival and be susceptible to BH3 mimetics targeting these proteins. In agreement with this latter point, the cells were also highly sensitive to low doses of PUMA BH3 peptide, which can only cause cytochrome c release by inhibiting the activity of pro-survival proteins and freeing any actively-bound pro-apoptotic BH3-only activator proteins to then activate BAX/BAK. Finally, we compared the level of overall apoptotic priming in patients with treatment-naïve disease versus those that are relapsed and found that treatment-naïve plasma cells are significantly more primed for apoptosis than those in patients at relapse (Figure 1B), which may contribute to increased therapy resistance in patients that have been previously treated. Overall, these results are consistent with other hematological disorders that have been analyzed in a similar manner^28,29,52^ and suggest that enhancing apoptosis may be an effective therapeutic strategy for controlling AL amyloidosis.

**Figure 1.**
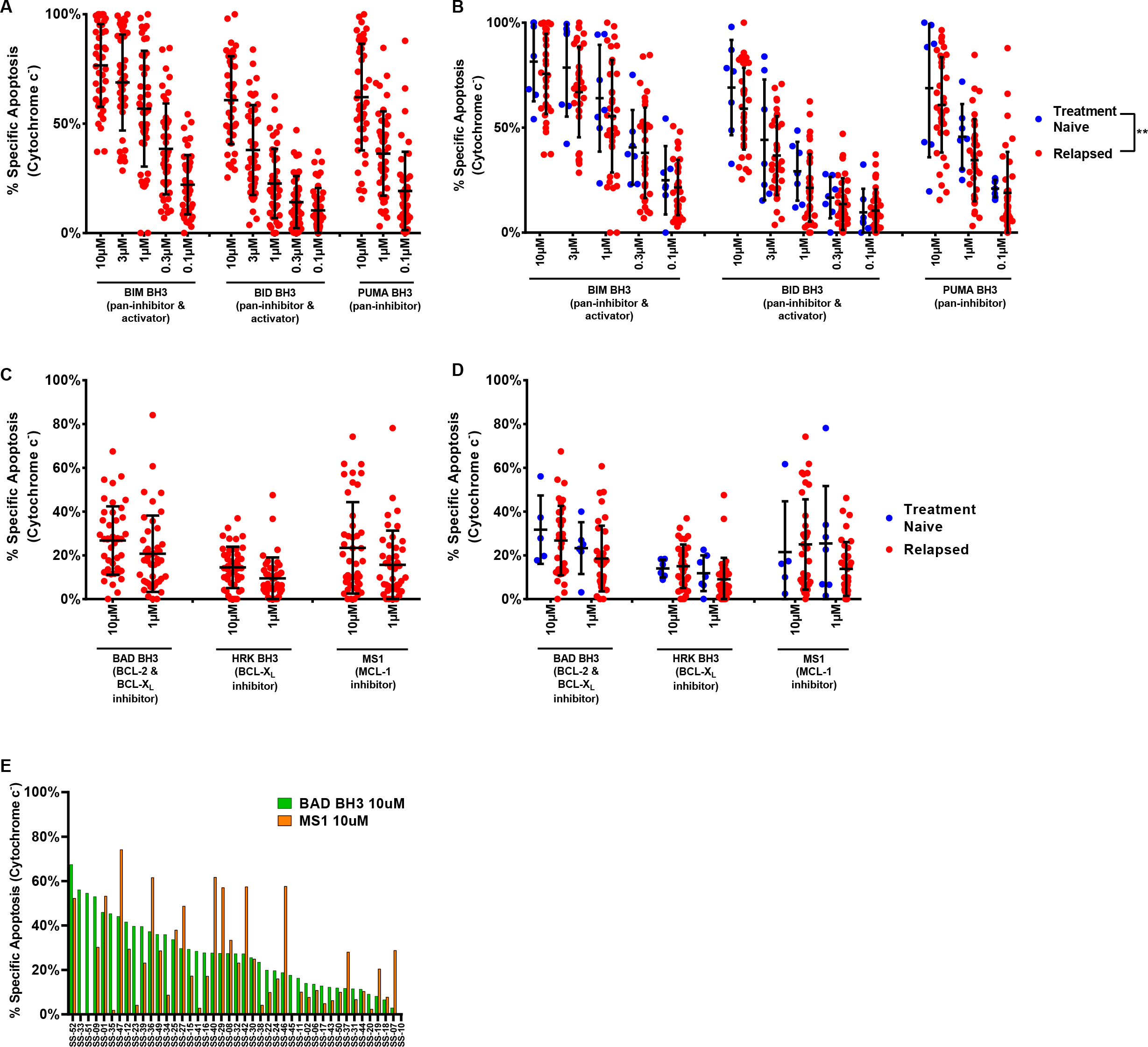
Plasma cells from patients with AL amyloidosis are primed for apoptosis. (A) Plasma cells were isolated from patient-derived bone marrow aspirates and BH3 profiled. Cells were exposed with pro-apoptotic BH3-only sensitizer peptides BIM, BID, and PUMA for 1 hour, then cytochrome c loss was measured by flow cytometry. (B) Patient samples were stratified into treatment naïve or relapsed subgroups according to their treatment status at the time of collection. (C) Plasma cells were BH3 profiled and exposed to peptides that inhibit pro-survival proteins to determine anti-apoptotic protein dependence. (D) Patient samples were stratified based on their treatment status at the time of collection. (E) Patients were ranked based on their response to BAD BH3 peptide treatment (BCL-2, BCL-X_L_ dependence) and their corresponding MS1 peptide response (MCL-1 dependence). Significance: ns, not significant; * *p* < 0.05; ** *p* < 0.01; *** *p* < 0.001; **** *p* < 0.0001.

We next sought to assess the extent of plasma cell dependence on pro-survival proteins by measuring MOMP in response to sensitizer BH3 peptides that each can only inhibit the activity of specific pro-survival BCL-2 family proteins. For this analysis, we exposed permeabilized plasma cells to the BAD BH3 (inhibits BCL-2, BCL-X_L_ and BCL-w), HRK BH3 (inhibits BCL-XL) and MS1 (inhibits MCL-1) peptides. Although responses were heterogeneous between patient samples, we observed that plasma cells released cytochrome c in response to the BAD BH3 and MS1 peptides, indicating dependence on BCL-2/BCL-XL or MCL-1, respectively. The responses to the HRK BH3 peptide were consistently lower than the other peptides, suggesting that BCL-XL is not a strong dependency for plasma cells and that the sensitivity to the BAD peptide is more an indication of BCL-2 dependence than BCL-XL. We also compared responses in treatment-naïve versus relapsed patients and found that plasma cells from relapsed patients tended to be less dependent on BCL-2 and more dependent on MCL-1 (Figure 1D), although the differences were not statistically significant. When examining the landscape of dependencies across samples (Figure 1E), we found that while some patients were sensitive to both BAD and MS1 (e.g. SS-52, SS-01), others were resistant to both (e.g. SS-10, SS-18), although the majority exhibited sensitivity to at least one peptide (e.g. SS-33, SS-51 for BAD; SS-07 for MS1). Taken together, these results suggest that inhibiting pro-survival proteins BCL-2 and/or MCL-1 will directly activate apoptosis in diseased plasma cells from patients with AL amyloidosis and enhance sensitivity to other apoptosis-inducing agents.

Although our BH3 profiling analysis allowed for a measurement of apoptotic dependencies without the need for ex vivo culture, we also sought to directly test the sensitivity of clonal plasma cells to BH3 mimetics. Whenever possible, a portion of ficolled mononuclear cells were apportioned for in vitro chemosensitivity assays using validated BH3 mimetics alone and in combination with agents that are currently used in treatment of this disease. After 24 hours of treatment in vitro, we found that plasma cells readily underwent apoptosis in response to the BCL-2 inhibitor ABT-199 (venetoclax), the BCL-2/BCL-XL inhibitor ABT-263 (navitoclax) and the MCL-1 inhibitor S63845 (Figure 2A), but were rarely sensitive to the BCL-X_L_ inhibitor WEHI-539, which was in agreement with our BH3 profiling data (Figure 1C). In addition, we again found that samples from treatment-naïve patients were more dependent on BCL-2 while relapsed patients were more dependent on MCL-1 (Figure 2B), suggesting that disease status may help determine which BH3 mimetic would be most efficacious for each patient.

**Figure 2.**
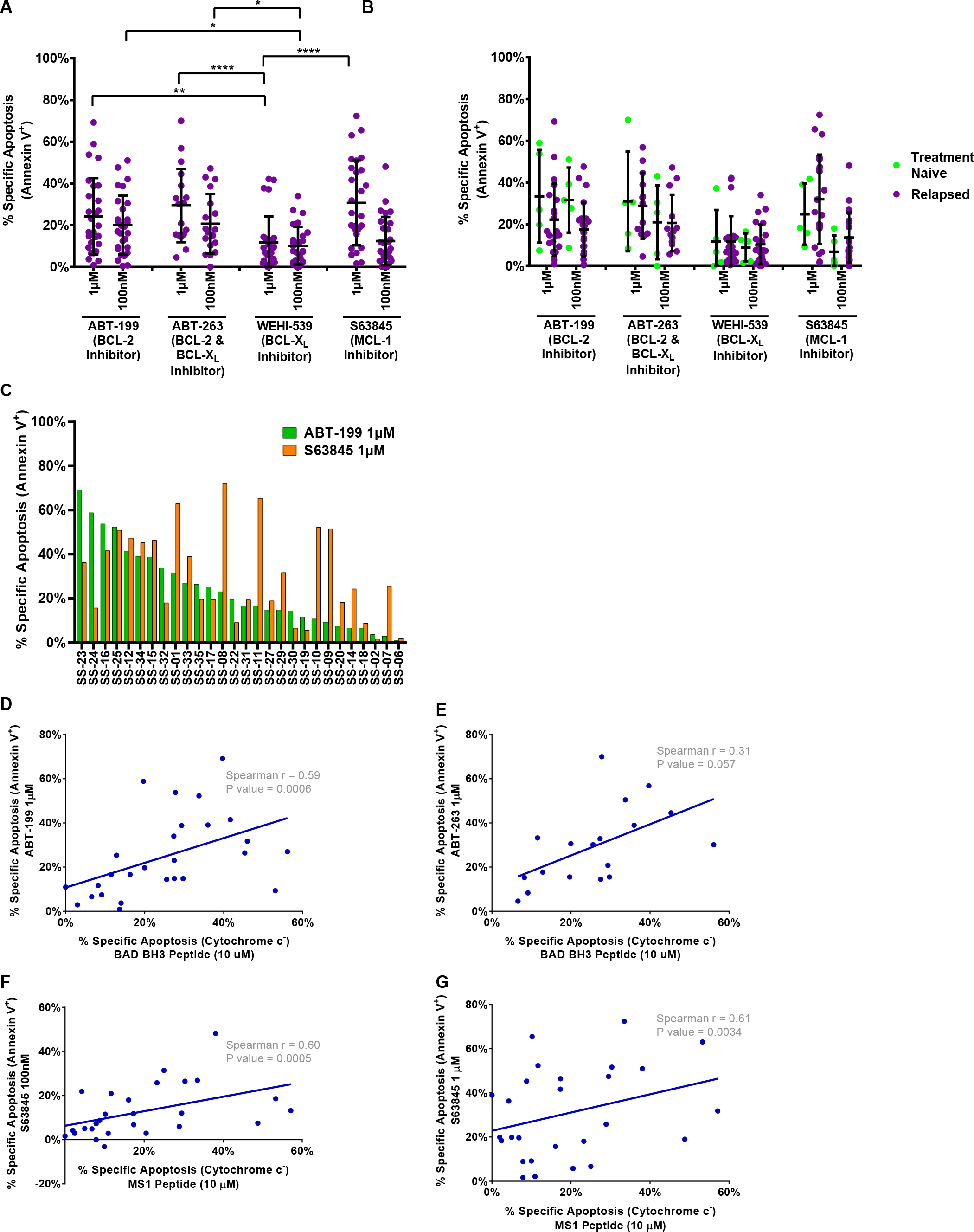
Plasma cells from patients diagnosed with AL amyloidosis undergo apoptosis in response to BH3 mimetics targeting BCL-2, MCL-1, and BCL-X_L_. BH3 profiling predicts ex vivo sensitivity to specific BH3 mimetics. (A) Plasma cells were isolated from patient bone marrow aspirates, cultured ex vivo, and treated with BH3 mimetics. Apoptosis was measured via Annexin V positivity after 24 hours. (B) Patient samples were stratified into treatment naïve or relapsed subgroups according to their treatment status at the time of collection. Patients’ apoptotic response to BH3 mimetics indicated. (C) Patients were ranked based on their apoptotic response to the ABT-199 (BCL-2 inhibitor) and their corresponding sensitivity to S63845 (MCL-1 inhibitor). The strength of correlation between clonal plasma cells’ apoptotic sensitivity via BH3 profiling at the time of isolation and their ex vivo chemosensitivity after 24 hour ex vivo treatment were measured for (D) BAD BH3 peptide treatment (indicating BCL-2, BCL-X_L_ dependence) and ABT-199 treatment (BCL-2 inhibitor), (E) BAD BH3 peptide treatment and ABT-263 treatment (BCL-2, BCL-X_L_ inhibitor), and MS-1 peptide treatment (MCL-1 dependence) and (F) 100 nM S63845 treatment (MCL-1 inhibitor) or (G) 1 µM S63845 treatment. Significance: ns, not significant; * *p* < 0.05; ** *p* < 0.01; *** *p* < 0.001; **** *p* < 0.0001.

The landscape of sensitivity to BH3 mimetics (Figure 2C) was similar to the results obtained via BH3 profiling, with patients exhibiting the same patterns of dependencies on pro-survival proteins. In fact, we found that in vitro sensitivity to BH3 mimetics was strongly correlated with sensitivity to the predicted peptides in the BH3 profiling analysis when comparing BAD BH3 response to ABT-199 or ABT-263 sensitivity (Figure 2D-E) and, to a lesser extent, when comparing MS1 peptide response to S63845 sensitivity (Figure 2F-G).

While we were encouraged by the sensitivity of plasma cells to BH3 mimetics as single agents, we also investigated the potential for combining these agents with standard of care therapies. Consistent with the strong activity of proteasome inhibitors as treatments for AL amyloidosis, we found that both bortezomib and ixazomib potently induced apoptosis in clonal plasma cells (Figure 3A-B). Further analysis demonstrated that sensitivity to proteasome inhibitors was highly correlated with cellular sensitivity to the BIM BH3 peptide in the BH3 profiling assay (Figure 3C-D), indicating that cells that are more primed for apoptosis tend to be more sensitive to these agents, which have been previously shown to induce apoptosis by enhancing ER stress, which increases pro-apoptotic signaling^53–55^. Importantly, combining proteasome inhibition with ABT-199 strongly enhanced apoptosis in clonal plasma cells, suggesting that this treatment combination may be effective clinically. However, adding S63845 to proteasome inhibition did not increase apoptosis in clonal plasma cells - this was unexpected based on the increased sensitivity to S63845 as compared to ABT-199 when used as single agents (Figure 2A). To determine the underlying mechanism that can be impairing additive induction of apoptosis, we BH3 profiled plasma cells during treatment with bortezomib to assess how the apoptosis pathway changes due to this stressor. We found that bortezomib strongly enhanced cellular sensitivity to the BAD BH3 peptide, indicating that dependence on BCL-2 is increased in these treated cells (Figure 3E). In comparison, there was only a mild difference in sensitivity to the HRK or MS1 peptides. These results suggested that expression of BCL-2 family proteins may change during bortezomib treatment, which has been previously demonstrated in neoplastic cells^53,56^. To test this, we measured the effects of bortezomib treatment on protein expression of key BCL-2 family members and found that, in all four patient samples examined, pro-apoptotic Noxa was consistently upregulated during proteasome treatment (Figure 3F-G) while treatment with thalidomide derivatives had no effect on expression of this protein. This accumulation of Noxa is likely due to the impairment of its normally short half-life^57^ and constant degradation via 26S proteosomes^58,59^. Noxa is an endogenous inhibitor of MCL-1, and the increased levels of this protein would effectively inhibit MCL-1 and shift cells toward apoptosis, thus causing increased dependence on other pro-survival proteins that are expressed (in this case being BCL-2). Thus, bortezomib treatment effectively mimics MCL-1 inhibition, thus potentially explaining the lack of increased apoptosis when combining MCL-1 inhibitors with proteasome inhibitors. We next tested the in vitro sensitivity of AL amyloidosis plasma cells to treatment with BH3 mimetics in combination with other standard therapies. ABT-199 again potently enhanced apoptosis in plasma cells when used in combination with thalidomide derivatives (Figure 4A-B) as well as dexamethasone (Figure 4C).

**Figure 3.**
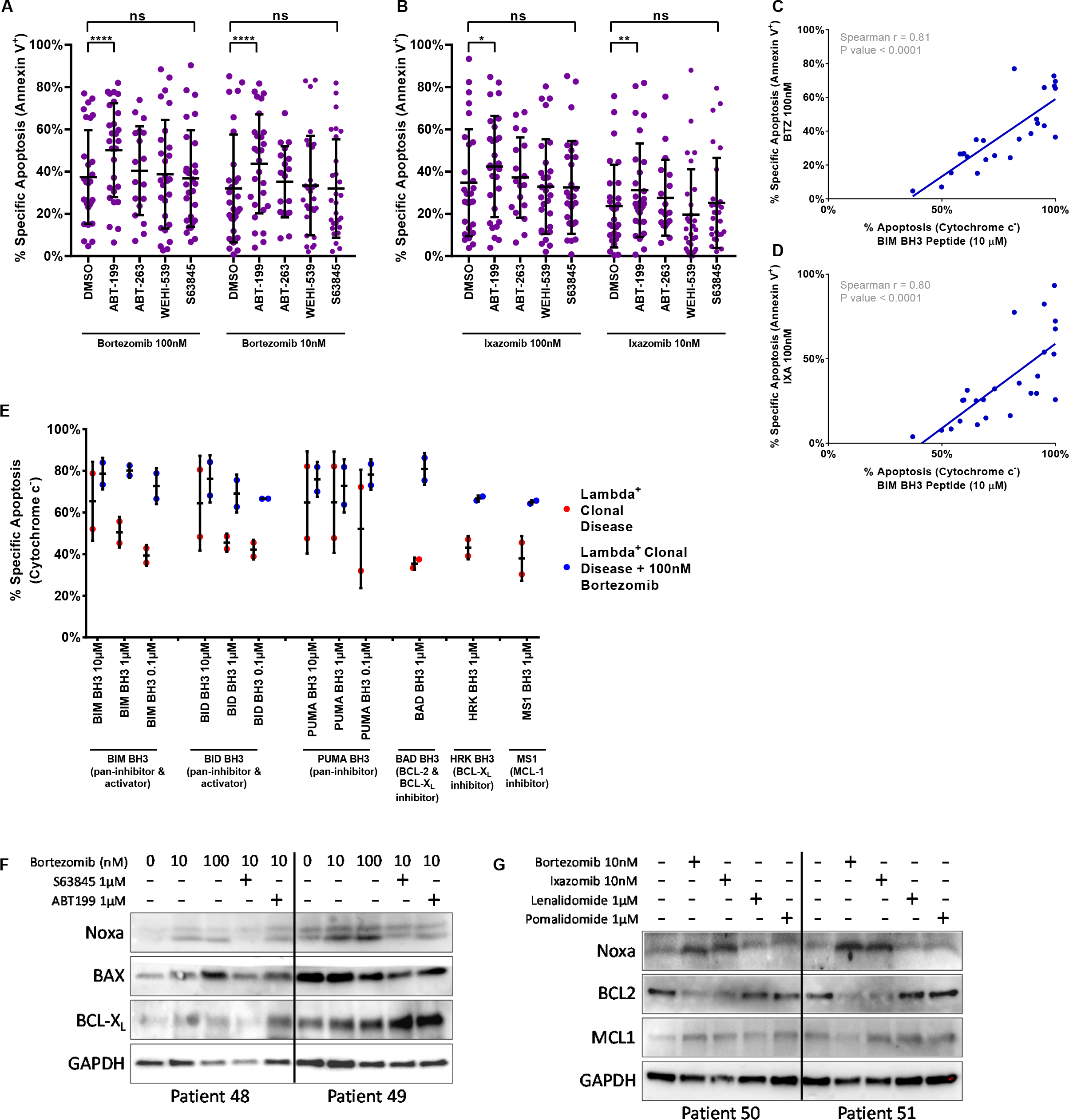
Plasma cells from patients with AL amyloidosis become more primed for apoptosis and more dependent on anti-apoptotic BCL-2 family proteins when treated with bortezomib, which was associated with increased Noxa expression. (A-B) Plasma cells were isolated from patient bone marrow aspirates, cultured ex vivo, and treated with (A) bortezomib and (B) ixazomib as single agents or in combination with BH3 mimetics. Apoptosis was measured via Annexin V positivity after 24 hours. Overall apoptotic priming of clonal plasma cells at the time of isolation was measured by sensitivity to the BIM BH3 peptide via BH3 profiling and correlated to the ex vivo sensitivity of the clonal plasma cell population in response to (C) bortezomib and (D) ixazomib treatments measured via Annexin V positivity after 24 hours. (E) Plasma cells were cultured ex vivo with or without 100 nM bortezomib and BH3 profiled after 24 hours. Lysates were prepared from patient-derived bone marrow mononuclear cells and western blot analysis was performed looking at changes in expression of BCL-2 family proteins in response to 24 hours of ex vivo culture with (F) bortezomib treatment as a single agent or in combination with ABT-199 and S632845. (G) Similarly, BCL-2 family protein expression was measured in response to 24 hour ex vivo treatment with bortezomib, ixazomib, lenalidomide, or pomalidomide. Significance: ns, not significant; * *p* < 0.05; ** *p* < 0.01; *** *p* < 0.001; **** *p* < 0.0001.

**Figure 4.**
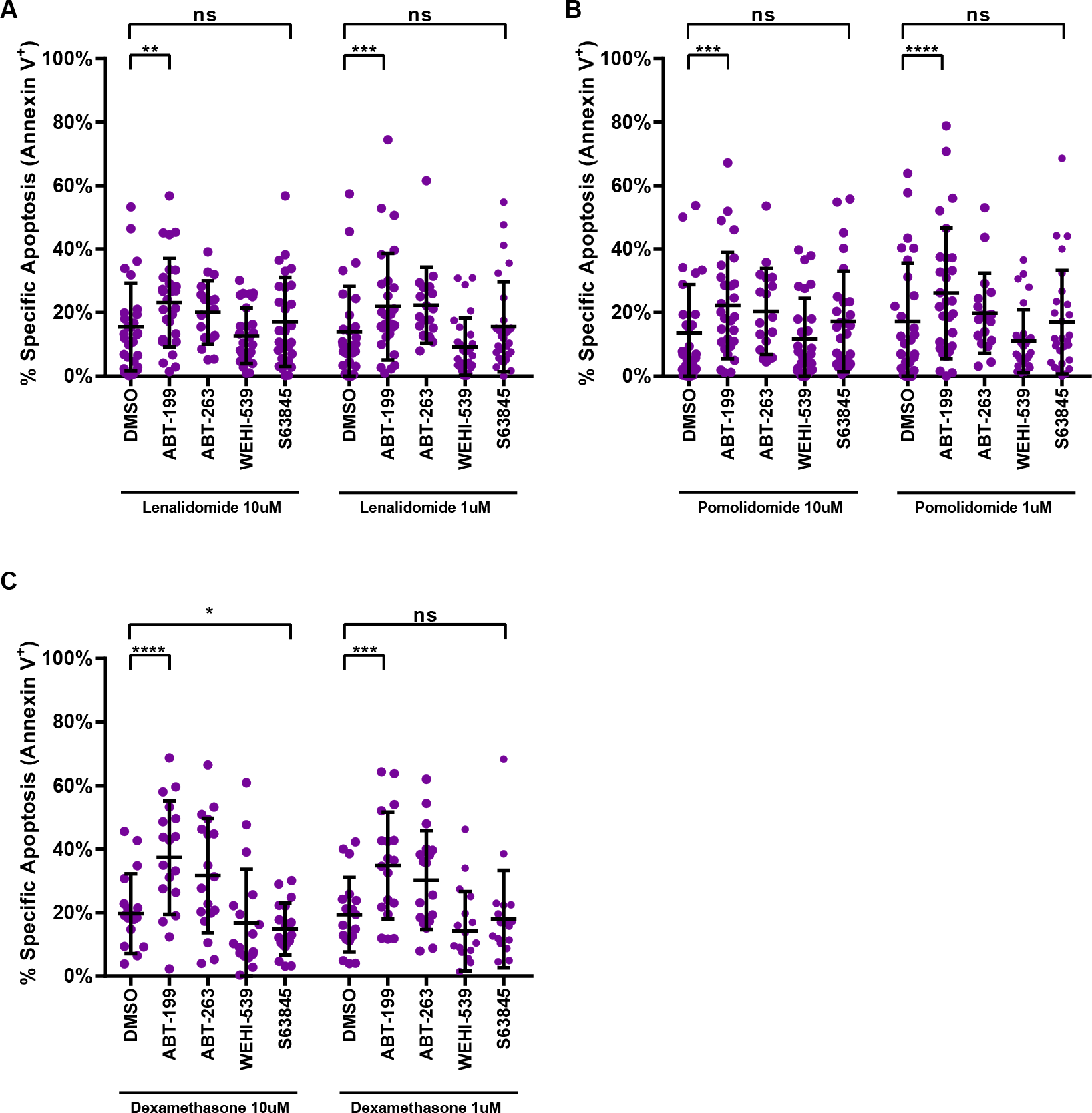
Combination treatment of lenalidomide, pomalidomide, or dexamethasone with ABT-199 results in increased plasma cell apoptosis. Plasma cells were isolated from patient bone marrow aspirates, cultured ex vivo, and treated with (A) lenalidomide, (B) pomalidomide, or (C) dexamethasone as single agents or in combination with BH3 mimetics. Significance: ns, not significant; * *p* < 0.05; ** *p* < 0.01; *** *p* < 0.001; **** *p* < 0.0001.

AL amyloidosis is typically diagnosed as either a κ or λ predominant disease and their clinical outcomes differ^60,61^. To better understand any differences in their regulation of BCL-2 family proteins, we stained for κ or λ light chains prior to BH3 profiling analysis to compare these two diseases. We found that clonal plasma cells from λ-predominant AL amyloidosis were significantly more primed for apoptosis than the clonal plasma cells from κ-predominant disease (Figure 5A). We further found that clonal λ-predominant plasma cells were also more responsive to sensitizer BH3 peptides, indicating that they are also significantly more dependent on BCL-2 (increased response to BAD BH3 peptide - Figure 5B) and tended to be more sensitive to the MCL-1 inhibiting MS1 peptide as well. The predominance of κ or λ may become a useful biomarker for potential assignment of therapy with BH3 mimetics clinically.

**Figure 5.**
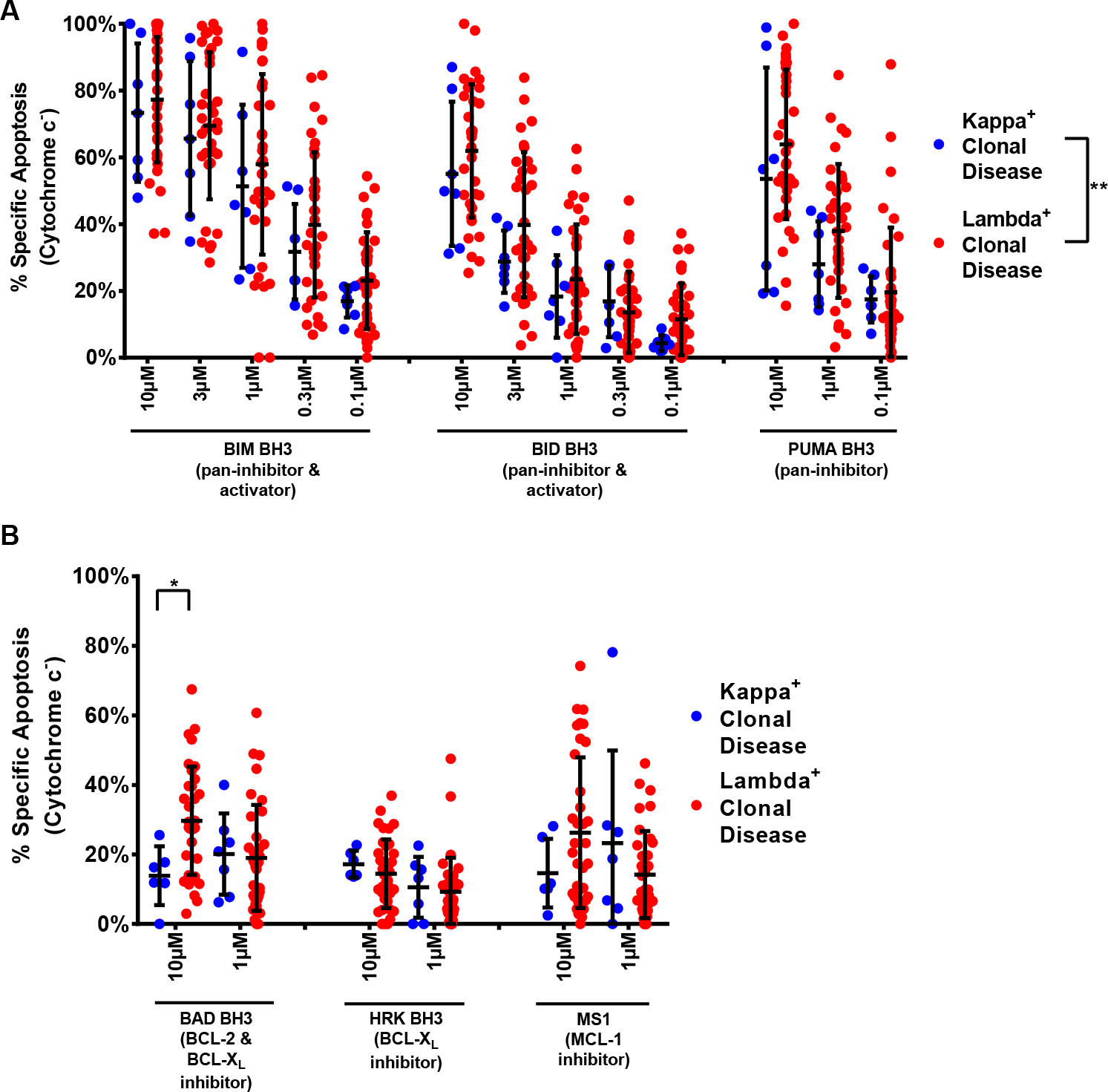
Clonal plasma cells from Lambda disease are more primed for apoptosis than clonal plasma cells from Kappa disease, more sensitive to BCL-2 inhibition. Clonal plasma cells isolated from Kappa- and Lambda-predominant AL amyloidosis were isolated and BH3 profiled to measure their (A) overall level of apoptotic priming via sensitivity to the BIM BH3, BID BH3, and PUMA BH3 peptides and their (B) dependencies on the pro-survival BCL-2 family proteins BCL-2, BCL-X_L_, and MCL-1. Significance: ns, not significant; * *p* < 0.05; ** *p* < 0.01; *** *p* < 0.001; **** *p* < 0.0001.

## Discussion

Herein, by studying apoptosis regulation directly in clonal plasma cells from patients diagnosed with AL amyloidosis utilizing several complementary approaches, we were able to make several key findings that may have important clinical implications. First, our BH3 profiling studies establish that clonal plasma cells have an intact and, in most cases, primed apoptosis pathway, as indicated by the high sensitivity of these cells to pro-apoptotic BIM and BID BH3 peptides. This high level of priming suggests that therapeutic interventions that can enhance apoptotic signaling are likely to be effective treatments in this disorder. Furthermore, we found that plasma cells are also highly sensitive to the pan-sensitizer peptide PUMA BH3, which can only inhibit pro-survival protein function but cannot directly active BAX or BAK^51,55,62^. This indicates that pro-survival proteins within these cells are actively sequestering pro-apoptotic proteins such as BIM, BID, BAX or BAK and effectively sets the stage for dependence on the function of pro-survival proteins and thus potential sensitivity to BH3 mimetics.

We also found that clonal plasma cells from treatment-naïve patients are more primed than those in relapsed disease, as we and others have previously described^29,35,52^. This suggests that chemotherapy treatments preferentially eradicate cells that are most highly primed for apoptosis, thus resulting in lower priming at the time of disease relapse due to the growth of more apoptosis-resistant, and thus likely more therapy-resistant, clonal plasma cells. We would thus expect that the highest likelihood of response in these patients with BH3 mimetics would be upon initial treatment.

In results from in vitro chemosensitivity, as well as BH3 profiling, assays we consistently found that clonal plasma cells in AL amyloidosis are highly dependent on both MCL-1 and BCL-2 for survival. Clinically, it may be possible to utilize a combination of both agents yet the potential for doing so and the toxicities that may be associated with this treatment are currently unknown. BCL-2 inhibitors alone are well tolerated with neutropenia being the primary toxicity^36,43,45^. MCL-1 inhibitors are being evaluated in early phase clinical trials and thus far seem to also be adequately tolerated but clinical data is still nascent. Although some studies have suggested that systemic MCL-1 inhibition may impair the function of the hematopoietic and cardiovascular systems^63^, this toxicity may be avoided by using sub-maximal dosing. In fact, studies have shown that even partial inhibition of MCL-1 can be strongly effective at controlling lymphoma growth in mouse models while being well tolerated^64^. Other studies have shown that most irreplaceable adult tissues are extremely resistant to apoptosis and are unlikely to be sensitive to MCL-1 inhibition^65^. Furthermore, MCL-1 has been reported to have important roles in maintaining mitochondrial homeostasis^66^ outside of its role as an apoptosis regulator. Thus, genetic deletion of MCL-1 may not be phenocopied by MCL-1 inhibition, especially since BH3 mimetics are specifically designed to inhibit the pro-survival activity of this protein. As data from clinical trials are reported, the extent of any toxicities and their optimal management will be better established.

To maximize the therapeutic potential of these apoptosis-sensitizing agents, BH3 mimetics may be used in combination with other drugs that induce apoptotic signaling, such as proteasome inhibitors. Interestingly, when we systematically combined BH3 mimetics with bortezomib or ixazomib, we found that the BCL-2 targeting agent ABT-199 most potently increased apoptosis in clonal plasma cells while combination with the MCL-1 inhibitor S63845 did not substantially enhance cell death. This contrasted sharply with the potent single-agent activity of S63845. We hypothesized that this may be due to changes in MCL-1 expression levels due to typical turnover of this ultra-short half-life protein^67^ being mediated by the target of bortezomib: the 26S proteasome. Although we did not observe any changes in expression of MCL-1, we found that all four of the patient samples that we tested exhibited an increase in the expression of Noxa during bortezomib or ixazomib treatment. Noxa is an endogenous specific inhibitor of MCL-1 and its overexpression would naturally inhibit the pro-survival activity of MCL-1. Noxa cannot bind to any other pro-survival proteins and thus targeting MCL-1 with S63845 in bortezomib-treated cells has no added benefit.

A prominent challenge to the deployment of BH3 mimetics is accurately assigning therapies to patients when heterogeneity in responses is expected. We found that BH3 profiling was a strong predictor of ex vivo sensitivity of clonal plasma cells to specific BH3 mimetics, suggesting that it may be an effective tool for personalizing therapy. This is further aided by our observation that bortezomib and ixazomib sensitivity was strongly correlated with overall apoptotic priming (response to the BIM BH3 peptide) suggesting that this assay may also help predict whether patients are likely to respond to proteasome inhibitors. We also found that λ-predominant AL amyloidosis exhibits higher levels of priming and dependence on both BCL-2 and MCL-1, suggesting that κ/λ status may also help assign therapy.

Historically, the lack of potent and specific BCL-2 family inhibitors has precluded the clinical evaluation of these agents for AL amyloidosis patients - this shortcoming has been recently overcome and agents targeting each of the pro-survival proteins are now in clinical trials for a variety of malignancies^40,44,68^. Our mechanistic and empirical data suggest that initial treatment of patients with λ-predominant AL amyloidosis with a proteasome inhibitor in combination with BCL-2 inhibitor would be highly efficacious and is likely to be well tolerated by patients, especially considering the acceptable safety profile seen for this combination in multiple myeloma^69^. For patients that may not tolerate bortezomib at diagnosis due to excessive disease-induced organ dysfunction, use of an MCL-1 inhibitor as a single agent may be effective at eliminating those cells that would typically be sensitive to bortezomib while being less toxic to healthy tissues affected by light chain deposition. If MCL-1 inhibition induced a reduction in clonal plasma cell populations and an improvement in organ function or patient performance status, initiation of combination therapy with a proteasome inhibitor and a BCL-2 inhibitor could then help eliminate remaining clonal plasma cells.

In summary, we have demonstrated that clonal plasma cell populations in AL amyloidosis demonstrate sensitivity to BH3 mimetics, and the sensitivity to treatment may potentially be predicted based on BH3 profiling or additional characteristics, such as κ/λ or prior treatment status. Although many BH3 mimetics are still under development, these may ultimately prove to be a well-tolerated and effective therapy that can be administered even in patients with AL amyloidosis at high risk for treatment toxicity and mortality. It is important to note that the insights we made within these studies are not only important for patients being treated for AL amyloidosis but also potentially other malignancies that have been reported to have apoptotic dependencies. These include lung adenocarcinomas^46,70^, leukemias^71,72^, lymphomas^73,74^, and even certain central nervous system tumors^75^. Our findings may therefore aid the effort to successfully deploy BH3 mimetics in AL amyloidosis, as well as multiple cancer types.

## Materials and methods

### Isolation of mononuclear cells

Bone marrow aspirates were collected from patients with treatment-naïve and relapsed AL amyloidosis under an Institutional Review Board-approved protocol at Boston Medical Center in Boston, MA. Samples ranging from 3-5 mL were diluted in PBS to 10 mL final volume. The diluted samples were then carefully layered over 4.5 mL of Ficoll-Paque PREMIUM (17-5442-02, GE Healthcare Biosciences, Uppsala, Sweden) and centrifuged at 450xg for 35 minutes. The layer of mononuclear cells was removed by pipette, added to PBS, and counted by hemocytometer.

### Flow cytometry-based BH3 Profiling

For each sample, 8×10^6^ mononuclear cells were isolated, centrifuged at 2000xg for 5 minutes, and resuspended in 150µL FACS Stain Buffer (2% FBS in PBS). Cells were then stained with the following conjugated cell-surface marker antibodies at 1:50 dilution: CD38 APC/Cy7 (102728, Biolegend, Dedham, MA) and CD138 Pacific Blue (356532, Biolegend). Cells were stained on ice for 25 minutes away from light. Cells were then centrifuged at 2000xg for 5 minutes and subjected to BH3 Profiling as previously described^49^. After BH3 profiling, cells were permeabilized for intra-cellular staining with a saponin-based buffer (1% saponin, 10% BSA in PBS) and intracellular-staining antibodies for Cytochrome C AlexaFluor 647 (612310, Biolegend), Kappa Light Chain FITC (11-9970-42, Invitrogen), and Lambda Light Chain (12-9990-42, Invitrogen), used at 1:2000, 1:50, and 1:50 dilutions, respectively. Cells were left to stain overnight at 4°C and analyzed by flow cytometry the following day. Cytochrome c positivity was measured on an Attune NxT flow cytometer.

### Ex vivo culture and viability measurement

Isolated mononuclear cells were plated at a concentration of 500,000 cells/mL in Dulbecco's Modified Eagle Medium (11995065, Gibco) with 10% Fetal Bovine Serum and 1% Penicillin/Streptomycin. Drugs were added to the culture as single agents and in combination treatments at the time of cell plating. Cells were harvested at 24 and 48 hours post treatment. After collection, cells were stained with the antibodies for CD38 and CD138 (see above). APC conjugated Annexin V antibody. APC conjugated Annexin V antibody was diluted 1:200 into 10x Annexin V binding buffer. Mixture was then added to the samples at 1/10^th^ of the volume of the sample and stained away from light for 20 minutes. Staining was fixed with 4% formaldehyde and 0.5% glutaraldehyde in 1x Annexin V binding buffer. After 10 minutes incubation, fixation was neutralized with N2 buffer (Tris Glycine). Cells were then stained for Kappa/Lamba light chains as described above. Analysis was completed on an Attune NxT flow cytometer.

### Statistical analysis

Two-way ANOVA (ordinary and Tukey's multiple comparisons test), Spearman correlation coefficient (r) and paired t tests were performed using the GraphPad Prism software (GraphPad Software). Significance: * *p* < 0.05; ** *p* < 0.01; *** *p* < 0.001; **** *p* < 0.0001.

