## Supplemental Figure and Table for "Clonal plasma cells in AL amyloidosis are dependent on pro-survival BCL-2 family proteins and sensitive to BH3 mimetics"

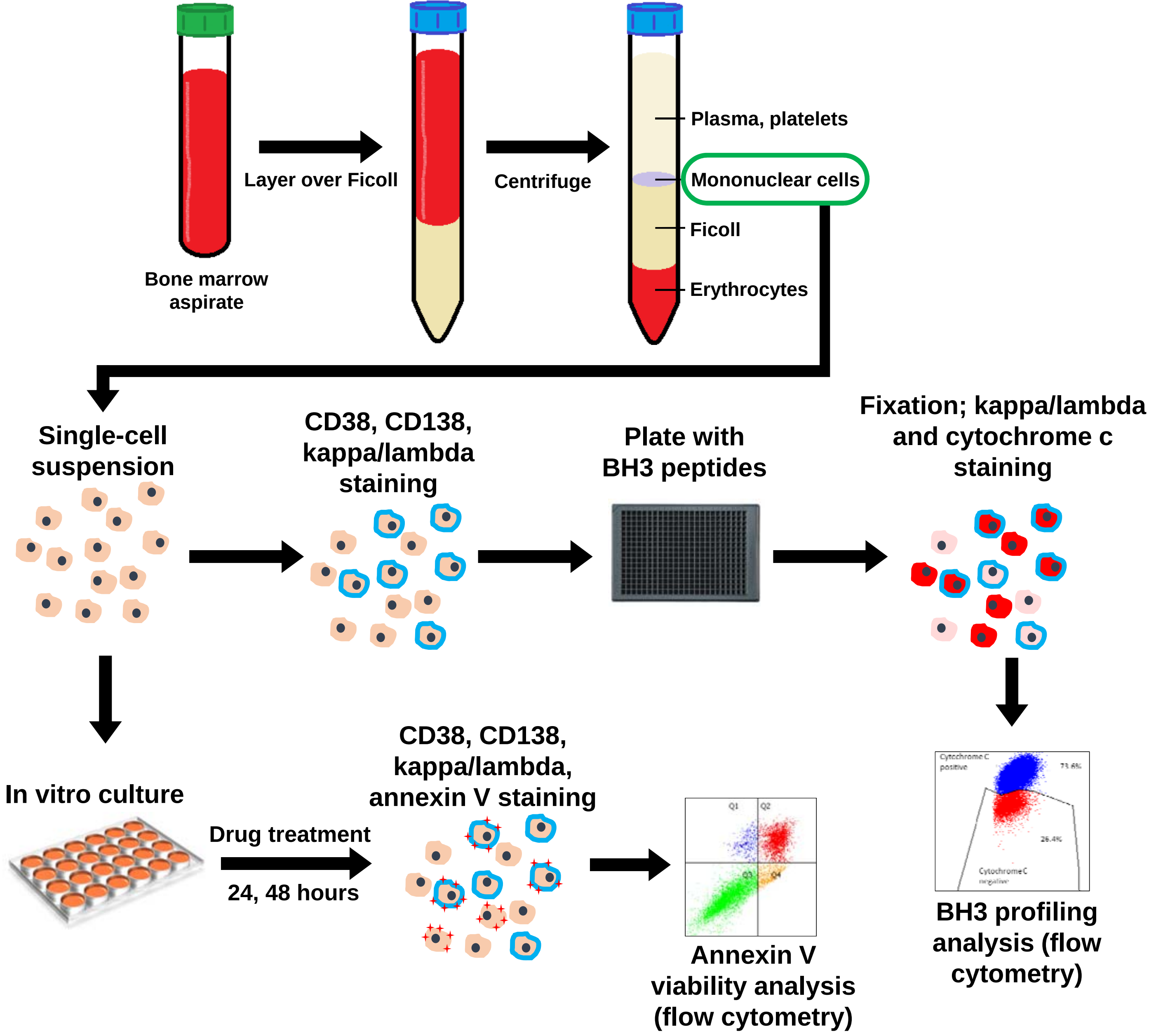


**Supplemental Figure 1: Processing of Primary AL amyloidosis samples for BH3 Profiling and in vitro chemosensitivity.**

| Patient # | t(11;14) | FISH | Organ involvement | Disease status | Prior treatments | Active therapy (if any) | Bone marrow report (% plasma cells and predominance based on IHC and ISH) | known type of amyloidosis | Disease status |
|---|---|---|---|---|---|---|---|---|---|
| **SS-01** | yes | gain 11, 13q14 | renal, cardiac, ANS | relapse | 2 cycle bor/dex, HDM/SCT | none | 5-10% plasma cells, lambda predominance | lambda | persistent disease |
| **SS-02** | yes | del 13, gain 11q | renal | PR | bor/dex | none | 10% plasma cells, lambda predominance | lambda | persistent disease |
| **SS-06** | no | 1q21 | cardiac, GI | VGPR | mel/dex | none | 5% plasma cells, no predominance | lambda | persistent disease (+IFE) |
| **SS-07** | unknown | unknown | renal | VGPR | 2 cycle bor/dex, HDM/SCT | none | 10-15% plasma cells, lambda predominance | lambda | persistent disease |
| **SS-08** | unknown | unknown | renal, ANS | VGPR | Ritux-VD, HDM/SCT, ritux maintenance | none | 5% plasma cells, lambda predominance | lambda | persistent disease |
| **SS-09** | no | normal | renal, ANS, cardiac | VGPR | CyBorD | none | 10% plasma cells, no predominance | lambda | persistent disease (+IFE) |
| **SS-10** | unknown | unknown | soft tissue | VGPR | CyBorD, HDM/SCT | none | 5-10% plasma cells, kappa predominance | kappa | persistent disease |
| **SS-11** | unknown | unknown | GI | PR | HDM/SCT x2, pom (6 days), bor/dex | none | 5-10% plasma cells, kappa predominance | kappa | persistent disease |
| **SS-12** | no | normal | renal | VGPR | CyBorD | none | 5-10% plasma cells, lambda predominance | lambda | persistent disease |
| **SS-14** | unknown | unknown | cardiac | VGPR | HDM/SCT | none | 10-15% plasma cells, lambda predominance | lambda | persistent disease |
| **SS-15** | no | 1q21 | renal | VGPR | RVD, bor | none | 5% plasma cells, lambda predominance | lambda | persistent disease |
| **SS-16** | no | normal | lymph node, lung | no prior therapy | no prior therapy | none | 10-15% plasma cells, lambda predominance | lambda | persistent disease |
| **SS-17** | unknown IGH rearrangement | 13q14, 14q32, -13 | renal, PNS, soft tissue, cardiac | CR | HDM/SCT | none | 15-20% plasma cells, lambda predominance | lambda | persistent disease |
| **SS-18** | yes | 14q32 | cardiac, renal, hepatic | PR | CyBorD(NEO), ixa/dex | ixa/dex | 10-15% plasma cells, lambda predominance | lambda | persistent disease |
| **SS-19** | unknown | unknown | cardiac | VGPR | bor/dex, rev/dex | none | 5-10% plasma cells, lambda predominance | lambda | persistent disease |
| **SS-20** | no | t914;20), +9, 1q21 | renal | no prior therapy | no prior therapy | none | unknown, the biopsy core was inadequate for eval | kappa | persistent disease (high sFLCs) |
| **SS-22** | unknown | unknown | renal, GI | no prior therapy | no prior therapy | none | 5-10% plasma cells, lambda predominance | lambda | persistent disease |
| **SS-23** | unknown | unknown | renal | VGPR | 2 cycle bor/dex, HDM/SCT, ixa/dex | none | 10% plasma cells, lambda predominance | lambda | persistent disease |
| **SS-24** | no | gain 9 and 15 | renal | no prior therapy | no prior therapy | none | 30-40% plasma cells, lambda predominance | lambda | persistent disease |
| **SS-25** | no | 11q13, 5' IGH del | renal | VGPR | HDM/SCT | none | 10% plasma cells, lambda predominance | lambda | persistent disease |
| **SS-27** | unknown | unknown | liver, renal, ANS | VGPR | CyBorD | none | 10-15% plasma cells, lambda predominance | lambda | persistent disease |
| **SS-29** | no | 1q21, -13, loss of IGH | Gi, cardiac, ANS | CR | CyBorD, HDM/SCT | none | 5-10% plasma cells, no predominance | lambda | persistent disease (+UIFE) |
| **SS-30** | unknown | unknown | renal | VGPR | bor/dex | bor/dex | 10-15% kappa plasma cells, kappa predominance | kappa | persistent disease |
| **SS-31** | unknown | unknown | renal, Factor X | CR | bor/dex | none | 5-10% plasma cells, lambda predominance | lambda | persistent disease |
| **SS-32** | unknown | unknown | renal | VGPR | HDM/SCT | none | 5-10% plasma cells, lambda predominance | lambda | persistent disease |
| **SS-33** | yes | 11 | lymph node, GI, cardiac, renal | no prior therapy | no prior therapy | none | 30-40% plasma cells, lambda predominance | lambda | persistent disease |
| **SS-34** | unknown | unknown | soft tissue, PNS, muscle, GI | **PR** | HDM/SCT, bor/dex, CyBorD, revl/dex | none | 10% plasma cells, lambda predominance | lambda | persistent disease |
| **SS-35** | yes | | cardiac | PR | CyBorD | CyBorD | 5% plasma cells, lambda predominance | lambda | persistent disease |
| **SS-36** | yes | | cardiac, renal | no prior therapy | no prior therapy | none | 5% plasma cells, no predominance | lambda | persistent disease (high sFLCs) |
| **SS-37** | unknown | unknown | renal, cardiac | CR | VMD, ritux/bor/dex, ritux maint | none | 5% plasma cells, kappa predominance | kappa | persistent disease |
| **SS-38** | unknown | unknown | liver, GI, renal, cardiac | VGPR | CyBorD, bor/dara/dex | bor/dara/dex | unable to interpret due to diffuse amyloid deposition | lambda | persistent disease (+UIFE) |
| **SS-39** | yes | -13 | GI, renal, cardiac, ANS | VGPR | mel/dex | none | 10-15% plasma cells, lambda predominance | lambda | persistent disease |
| **SS-40** | yes | loss 13q14, loss 13q34, 5' del, gain 9,11,and 15 | GI | PR | CyBorD+dara | CyBorD+dara | 5% plasma cells, no predominance | lambda | persistent disease (+IFE) |
| **SS-41** | unknown | unknown | GI, cardiac | PD | rev/mel/dex, pom/dex, ixa/dex, dara | none | 10-15% plasma cells, lambda predominance | lambda | persistent disease |
| **SS-42** | no | normal | renal, cardiac | VGPR | bor/dex, bor/dex/ritux, cytoxan/bor/dex/ritux | Cytoxan/ritux/dex | 10% plasma cells, lambda predominance | lambda | persistent disease |
| **SS-43** | no | 1q21, -13, del 3' | renal, cardiac | VGPR | CyBorD | none | 5-10% plasma cells, no predominance | lambda | persistent disease (+IFE) |
| **SS-44** | unknown | unknown | soft tissue, cardiac, renal | VGPR | HDM/SCT, Rev/dex | none | 10% plasma cells, lambda predominance | lambda | persistent disease |
| **SS-45** | | 11 | **soft tissue** | **PR** | **none** | **none** | 5% plasma cells, kappa predominance | kappa | persistent disease |
| **SS-46** | yes | gain 1 and 11 | cardiac, GI, soft tissue | PR | bor/dex | none | 10% plasma cells, no predominance | lambda | persistent disease (sFLCS, +IFE) |
| **SS-47** | unknown | unknown | renal | VGPR | HDM/SCT | none | 10-15% plasma cells, lambda predominance | lambda | persistent disease |
| **SS-49** | no | normal | cardiac, renal | CR | bor/dex | none | 5-10% plasma cells, no predominance | lambda | persistent disease (+IFE) |
| **SS-50** | yes | | renal | VGPR | CyBorD | none | 5% plasma cells, no predominance | kappa | persistent disease (sFLC ratio) |
| **SS-51** | no | gain 1q21, 3' del | renal, cardiac | PR | BorD, HDM/SCT | none | 15% plasma cells, lambda predominance | lambda | persistent disease |
| **SS-52** | unknown | unknown | GI, renal | PR | HDM/SCT, CyBorD, Bor/dex | bor/dex | 5% plasma cells, lambda predominance | lambda | persistent disease |
| **SS-53** | unknown | unknown | nasopharyngeal/larynx | N/A | no prior therapy | none | 5% plasma cells, no predominance | kappa | persistent disease |

**Supplemental Table 1: Patient characteristics.**
